# Assessment of the efficacy of field and laboratory methods for the detection of *Tropilaelaps spp*

**DOI:** 10.1101/2024.03.26.586849

**Authors:** Maggie C. Gill, Bajaree Chuttong, Paul Davies, Dan Etheridge, Lakkhika Panyaraksa, Victoria Tomkies, George Tonge, Giles E. Budge

## Abstract

*Tropilaelaps* spp. are invasive mites that cause severe disease in *Apis mellifera* colonies. The UK has deployed an elaborate surveillance system that seeks to detect these mites early in any invasion to allow the best opportunity to eradicate any incursion. Effective field and laboratory protocols, capable of reliably detecting low numbers of mites, are key to the success of any intervention. Here we compared the efficacy of established field monitoring using brood removal with an uncapping fork, and brood ‘bump’ methods with novel methods for *Tropilaelaps* detection modified from *Varroa* monitoring schemes. In addition, we monitored the efficacy of the laboratory method for screening for mites in hive debris by floating mites in ethanol. Our results clearly indicated that novel methods such as uncapping infested brood with tweezers, catching mite drop using sticky traps and rolling adult bees in icing sugar were all significantly more likely to detect *Tropilaelaps* than existing methods such using an uncapping fork on infested brood, or the brood ‘bump’ method. Existing laboratory protocols that sieved hive debris and then floated the mite containing layer failed to detect *Tropilaelaps* mites and new efficacious protocols were developed. Our results demonstrated that the national surveillance protocols for *Tropilaelaps* mite detection required modification to improve the early detection of this damaging invasive mite.

## Introduction

*Tropilaelaps* mites are native brood parasites of the Asian honey bee species *Apis dorsata, Apis breviligula* and *Apis laboriosa* which have successfully jumped host to introduced colonies of *Apis mellifera*. The current range of *Tropilaelaps* mites is not fully characterised (Brown. *et al*, 2002), but current reports restrict these mites to Asia and bordering areas (Anderson & Roberts, 2013; Aydin, 2022). What is clear is that infestations of *A. mellifera* by *Tropilaelaps mercedesae* occur in regions well beyond the distribution of their native Asian honey bee hosts, making the mite an emerging global threat to *A. mellifera* that has the potential to parallel the near-global spread of *Varroa destructor* (Anderson and Morgan 2007; Ramsey, 2021; Thompson, *et al*. 2002).

*Tropilaelaps spp*. have a shorter phoretic phase, a faster rate of reproduction than *Varroa* and have the ability to mate outside brood cells (Chantawannakul, *et al*. 2018). These life history traits allow populations of *Tropilaelaps spp*. to increase more rapidly than *Varroa*. Brood feeding causes physical damage (Khongphinitbunjong, *et al*. 2015) and *Tropilaelaps* mites are known to transmit viruses which have the potential to kill or shorten the life of the developing bee (Dainat *et al*. 2008; Forsgren *et al*. 2009; Khongphinitbunjong *et al*. 2015). Natural behaviours such as frequent absconding, migration and high levels of grooming and hygienic behaviour of the natural Asian honey bee hosts play an important role in the control of *Tropilaelaps spp*. (Koeniger *et al*. 2002), resulting in low infestation levels and concomitant low colony mortality (Buawangpong *et al*. 2013). *A. mellifera* has not yet evolved a symbiotic parasite-host relationship, resulting in high mite levels and high rates of colony mortality (Phokasem, *et al*. 2019).

The presence of brood is thought to be a prerequisite for *Tropilaelaps* mites to survive within an *A. mellifera* colony, with *T. clareae* demonstrating survival of only three days in artificial conditions (Tran, *et al*. 1993; Sharma, *et al*. 1998). There is suspicion that, under natural conditions, *Tropilaelaps spp*. could survive for longer periods in the absence of brood (Brown, *et al*. 2002), and *T. mercedesae* has been shown to overwinter in *A. mellifera* colonies in temperate climates where brood is often absent (Truong, *et al*. 2021).

Increased global trade and changes in beekeeping practices provide transmission routes for *Tropilaelaps spp*. which include live honey bee sales, migratory beekeeping and the potential distribution of feral colonies or swarms on shipping containers and cargo vessels (EFSA, 2013). *Tropilaelaps spp*. are statutory notifiable pests in most countries worldwide because of a combination of likely global distribution coupled with significant economic losses because of high mortality rates in *A. mellifera* colonies (APHA, 2017). The National Bee Unit (NBU) is a government body responsible for the surveillance of honey bee (*A. mellifera*) colonies in England and Wales. Their primary obligation is surveillance for the presence of the statutory notifiable bacterial diseases American foulbrood (caused by *Paenibacillus larvae*) and European foulbrood (caused by *Mellissococcus plutonius*) plus the statutory notifiable pests Small Hive Beetle (*Aethina tumida*) and *Tropilaelaps spp* (Brown *et al* 2014). The early interception of exotic pest incursions is imperative to increase the chances for the successful eradication of invasive pest species (Keeling *et al*. 2017). The UK operates an extensive monitoring programme for *Tropilaelaps spp*. that includes exotic pest risk points that are designated using a risk based classification system, with ports and queen importers identified as the highest incursion risk of an exotic pest incursion. A network of enhanced sentinel apiaries (ESA) are positioned in close proximity to the exotic pest risk points where bee inspectors from the NBU carry out colony inspections specifically targeted at the detection of *Tropilaelaps spp*. ESAs are complemented by voluntary sentinel apiaries (VSA), where volunteer beekeepers close to an exotic pest risk point regularly submit floor debris samples for laboratory analysis for *Tropilaelaps spp*. with the risk analysis and prioritisation of exotic pest risk points being based on the work of Keeling *et al*. (2017).

The NBU operate a standard operating procedure (SOP) for the detection of *Tropilaelaps spp*. in sentinel apiaries (ESA and VSA) based on techniques described by Pettis *et al*. (2013) to include three surveillance techniques:

1. The removal and inspection of 200 cells of sealed brood using an uncapping fork;
2. The ‘bump method’, where brood frames are hit against a solid surface to dislodge mites onto a white monitoring tray;
3. The laboratory screening of hive floor debris samples for *Tropilaelaps spp* (and *A. tumida*);

Should *Tropilaelaps* be detected using the above three methods, then enhanced surveillance would be completed using a miticide in conjunction with a sticky floor insert. It is worth noting that beekeepers often object to the damage caused by the bump method as well as the destructive nature of brood uncapping.

The aim of this study was to determine the efficacy of the existing field and laboratory protocols employed for the detection of *Tropilaelaps spp*. as part of the national exotic pest surveillance scheme and test novel methods for mite detection to improve the chance of national surveillance systems to detect these damaging mites early in any invasion.

## Methods

### Field inspections

In total, 60 colonies of *A. mellifera* were sourced from six apiaries in Chiang Mai province Thailand (Figure.1). Each colony was assessed using all 4 of the field protocols detailed below. Only one method for monitoring mite drop could be deployed on each colony, and so colonies were randomly allocated to receive either oil or sticky inserts, as both these media are known to trap mites (De Guzman, *et al*. 2017). Beekeepers were asked to not treat colonies for mites for at least two brood cycles prior to assessment, but colony management was not controlled prior to this point. Multiple detection methods were trialled, and the numbers of *Tropilaelaps* were counted for each method on each colony.

**Figure 1.**
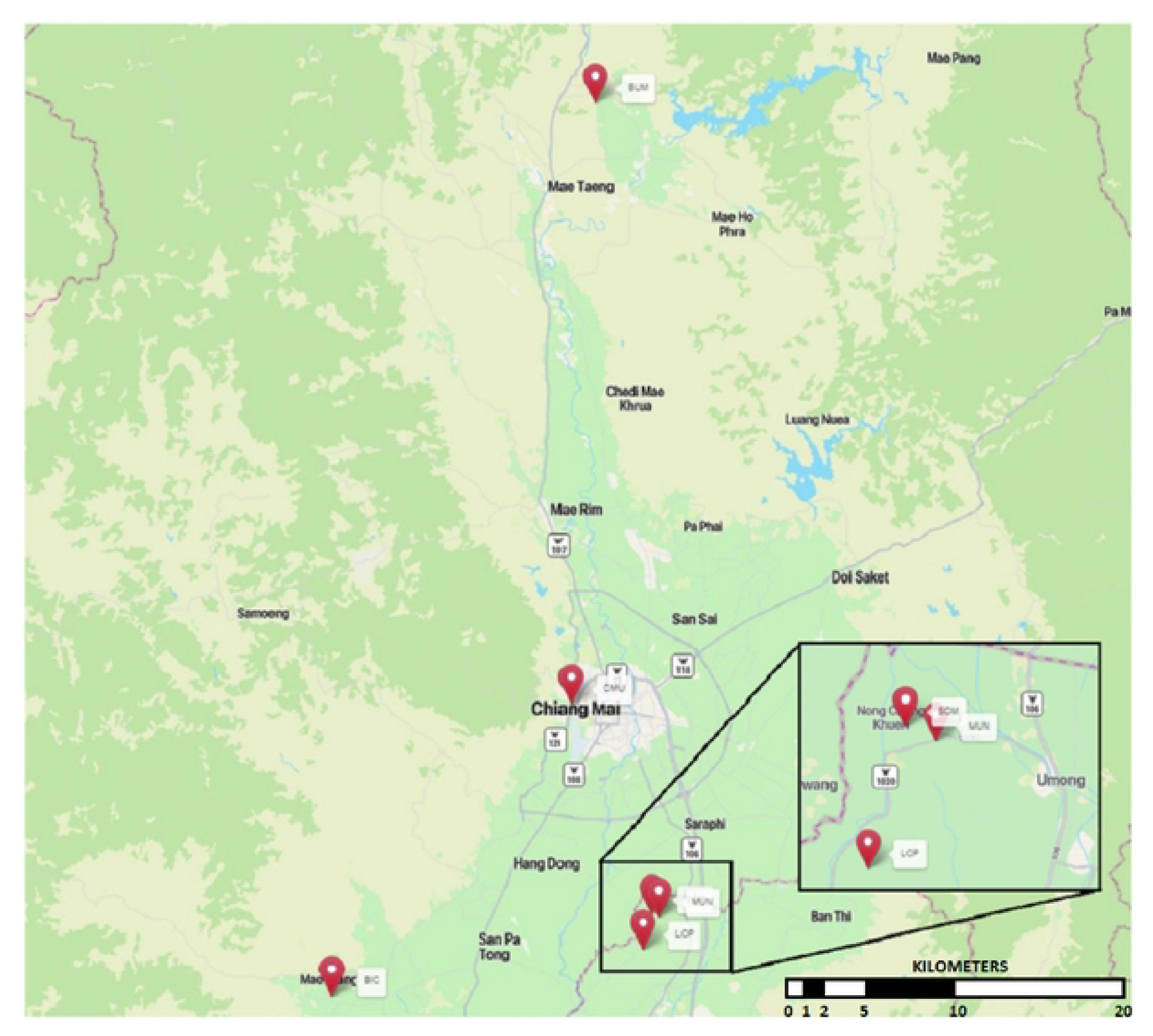
Map of apiary locations in Thailand.

### Field protocols

#### 1. Brood uncapping

The SOP for the detection of *Tropilaelaps spp*. specified that 200 sealed drone brood cells containing pink eyed pupae should be uncapped using an uncapping fork, however these were rarely present in the sampled colonies. When carried out mites became trapped by exudate from the damaged pupae or were difficult to identify amongst the exudate from the damaged pupae. As such, the method was modified to sample a 10 × 10 rhombus template of 100 sealed brood cells. Fine-nosed forceps were used to remove the immature honey bee from each cell and the larva or pupae were carefully inspected for mites. If brood from the final larval instar (day 9) to the pharate pupal stage (day 13) or brood that was post pink eyed stage (day 18) were removed, then *Tropilaelaps* mites tended to not be found on the pupae, but instead remained in the cells. They could be dispersed by gently blowing over the brood or by using a fine artist brush dipped in honey to sweep the cells. Typically, *Tropilaelaps* mites would be seen on extracted pupae that were between 14 to 17 days old. A headtorch and an illuminated 40x magnification hand lens was also employed to aid mite discovery.

#### 2. Comb bump method

The current surveillance procedure was adapted from the bump method described by Pettis *et al*. (2013). The SOP originally specified that adult bees were shaken from every brood frame and then each brood frame was held horizontally over a white tray with the wooden edge of the frame firmly hit twice per side on the edge of the tray allowing mites to fall into the tray. Beekeepers expressed concerns about the damage this caused the brood, therefore this method was refined, and adult bees were removed from one sealed and one unsealed brood frame selected at random. Each frame was held horizontally over a white tray with the wooden edge of the frame firmly hit four times per side on the edge of the tray allowing mites to fall into the tray. The tray was then examined for the presence of mites with a hand lens (x40) and head torch.

#### 3. Hive debris

Hive floor debris from open mesh floors are routinely collected from colonies and sent to the laboratory for analysis following a set protocol. Thailand beekeepers use an adapted Langstroth hive where hive floors are attached to the main hive brood body. As such, all the brood frames had to be removed to access the floor debris and samples were bagged in ziplock bags for later laboratory analysis.

In the UK, sticky inserts would normally be deployed on *Varroa* floors, but the configuration of Thai hives meant that floor inserts had to be modified slightly. Floor inserts were made up of a shallow wooden frame supporting a plastic mesh covering. A commercially available sticky insert (Vita Europe) was used in half the colonies and a laminated card coated with a thin layer of vegetable oil was used in the remaining colonies. No acaricide strips were used to promote mite drop as specified in the SOP as experimental permits could not be obtained to use UK approved acaricide strips. As such, only natural mite drop was recorded. Floor inserts were inserted for 24 hours, and mite counts completed in the laboratory.

Laboratory mite surveillance utilised methods for detection of *Tropilaelaps spp*. in floor debris samples outlined by Pettis *et al*. (2013), Anderson *et al*. (2013) and laboratory methods optimised in the National Bee Unit diagnostic laboratory. To assess the presence of mites, Endecott sieves were stacked on top of each other in decreasing size (1.7 mm, 1 mm, 0.7 mm and 0.355 mm) and hive debris placed in the top sieve. The sample was then flushed through the sieves using tap water passed through a small shower head, resulting in size separation of the debris. The smallest sieve contents (0.355 mm) were placed in a small washing up bowl of demineralised water, allowing mites to float. The sample was assessed in good light, using either a large magnifying glass or a jeweller’s hand lens (x40).

#### 4. Monitoring mites on adult bees

The frame containing the queen was found and set to one side. The adult bees from one frame of mostly open brood and one frame of mostly closed brood were shaken into a smooth-sided bowl. Approximately 300 adult bees were scooped into each of three *Varroa* EasyCheck mite counters (Veto Pharma). The first counter was treated with 300 ml of ethanol to dislodge the mites, and the mites counted through the clear base after one minute of swirling, essentially as described by Pettis *et al*. (2013). The second counter was treated with 25 g of icing sugar before the bees were rolled for one minute. The *Varroa* EasyCheck counter was then left to stand out of direct sunlight for one minute before the icing sugar was shaken out onto a white tray and assessed for the presence of mites. This process was repeated after a further two minutes to allow the bees further opportunity to groom. The third counter was treated with a steady supply of CO_2_ until the bees were fully anaesthetised. The bees were then gently rolled for fifteen seconds, before being shaken over a white tray and checked for the presence of mites using a hand lens (x40) and headtorch.

### Statistical analyses

To prevent bias towards monitoring method due to variation in mite levels between colonies, colonies were only included where data were gathered for all monitoring methods. All models were implemented using R version 4.2.3 (R Core team, 2018). First, we used a binomial generalised linear model (GLM) with a logit link function, with the presence of mites as a response variable and monitoring method and apiary as the independent variables. Second, we used zero-inflated regression models for count data via maximum likelihood implemented in the *pcsl* package (Zeileis, *et al*. 2008), with the number of mites as the response variable and monitoring method and apiary as the independent variables. Model fit was compared using the Akaike Information Criterion (AIC) and 95% confidence intervals of the estimates were calculated using the coefficients and standard error from the GLM summary table.

The data were partitioned into three different sets to account for the fact that half the colonies had mite drop monitored using oil, and the other half used sticky traps. First, the method of mite drop was removed to retain the highest number of observations for the remaining monitoring treatments. Second, the data were filtered to only those colonies with data using oil to monitor mite drop, and the third partition contained only those colonies with data using sticky traps to monitor mite drop. The efficacy of the monitoring methods was expressed as the odds ratio of detecting mites compared to washing mites off 300 adult bees with ethanol as the standard detection method. Finally, we investigated competition between *Varroa* and *Tropilaelaps* mites using a binomial generalised linear model with a logit link function, with the presence of *Varroa* mites as a response variable with monitoring method and *Tropilaelaps* presence as independent variables.

## Results

Of the 60 colonies that were examined, colony size averaged 5.3 frames of bees (±5.1) and 3.8 (±2.7) frames of brood. *T. mercedesae* was detected in 80% of colonies and 100% of apiaries. In total, seven colonies were removed from the analysis because the results from one or more monitoring method was absent. Of these seven colonies, five were deemed too small to provide the necessary brood samples for monitoring, and the uncapping method was incorrectly deployed in two colonies. In total, 53 colonies were assessed for the efficacy of alcohol, brood bump, CO_2_, uncapping and sugar roll. Mite drop was monitored using oil in 24 colonies and using sticky traps in 29 colonies.

### Comparing the ability of each method to detect Tropilaelaps mites

The AIC was consistently lower for the binomial GLM containing treatment and apiary (All colonies = 278.55; Oil drop only = 160.52; Sticky drop only = 193.24), compared to treatment alone (All colonies = 301.78; Oil drop only = 180.67; Sticky drop only = 203.61), and so this was chosen as our final model. Method of mite detection had a significant effect on the presence of *Tropilaelaps* mites in the colony for all three analyses (Table 1).

**Table 1.** Summary of binomial GLMs comparing *Tropilaelaps* mite detection using all methods in the absence of mite drop (All colonies; n = 53); including only those colonies that received oil to monitor mite drop (Oil drop only; n = 24); and including only those colonies that received sticky traps to monitor mite drop (Sticky drop only; n = 29).

| All colonies | Method/apiary | Estimate | St.error | Z value | P value |
| --- | --- | --- | --- | --- | --- |
|  | (Intercept) | -1.11 | 0.469 | -2.37 | <0.05 |
| *Method* | Bump | 0.23 | 0.477 | 0.48 | NS |
|  | CO<sub>2</sub> | -0.71 | 0.542 | -1.3 | NS |
|  | Uncapping | 2.79 | 0.516 | 5.41 | <0.001 |
|  | Sugar roll | 1.47 | 0.464 | 3.17 | <0.01 |
| *Apiary* | Bum | 0.60 | 0.445 | 1.36 | NS |
|  | CMU | -1.85 | 0.597 | -3.09 | <0.01 |
|  | Lop | -1.46 | 0.640 | -2.28 | <0.05 |
|  | Mun | -0.64 | 0.557 | -1.16 | NS |
|  | Som | 0.35 | 0.529 | 0.66 | NS |
| **Oil drop only** | (Intercept) | -1.08 | 0.662 | -1.63 | NS |
| *Method* | Bump | 0 | 0.703 | 0 | NS |
|  | CO<sub>2</sub> | -1.22 | 0.818 | -1.49 | NS |
|  | Uncapping | 2.34 | 0.753 | 3.11 | <0.01 |
|  | Oil drop | 0.24 | 0.694 | 0.35 | NS |
|  | Sugar roll | 1.36 | 0.698 | 1.95 | ?? |
| *Apiary* | Bum | 1.27 | 0.605 | 2.1 | <0.05 |
|  | CMU | -2.29 | 0.935 | -2.45 | <0.05 |
|  | Lop | -0.57 | 0.765 | -0.75 | NS |
|  | Mun | -0.95 | 0.799 | -1.18 | NS |
|  | Som | 0.66 | 0.719 | 0.91 | NS |
| **Sticky drop only** | (Intercept) | -1.6 | 0.645 | -2.47 | <0.05 |
| *Method* | Bump | 0.45 | 0.679 | 0.67 | NS |
|  | CO<sub>2</sub> | -0.28 | 0.749 | -0.37 | NS |
|  | Uncapping | 3.4 | 0.75 | 4.53 | <0.001 |
|  | Sticky drop | 1.51 | 0.65 | 2.32 | <0.05 |
|  | Sugar roll | 1.67 | 0.651 | 2.57 | <0.05 |
| *Apiary* | Bum | 0.32 | 0.55 | 0.59 | NS |
|  | CMU | -1.33 | 0.694 | -1.92 | NS |
|  | Lop | -2.22 | 0.961 | -2.31 | <0.05 |
|  | Mun | 0.32 | 0.644 | 0.5 | NS |
|  | Som | 0.77 | 0.642 | 1.19 | NS |

When comparing detection methods across all 53 colonies, it was clear that both uncapping brood and a sugar roll of bees were more likely to detect *Tropilaelaps* mites than floating the floor debris in alcohol. Bumping the brood and using CO_2_ to knock down the mites on adult bees were not better than alcohol (Table 1; Figure 2A). When restricting to those colonies where mite drop was monitored using oil, oil offered no significant improvement over alcohol (Table 1; Figure 2B). When restricting to those colonies where mite drop was monitored using sticky traps, the sticky traps were significantly more likely to detect the presence of *Tropilaelaps* mites when compared to alcohol (Table 1; Figure 2C).

**Figure 2.**
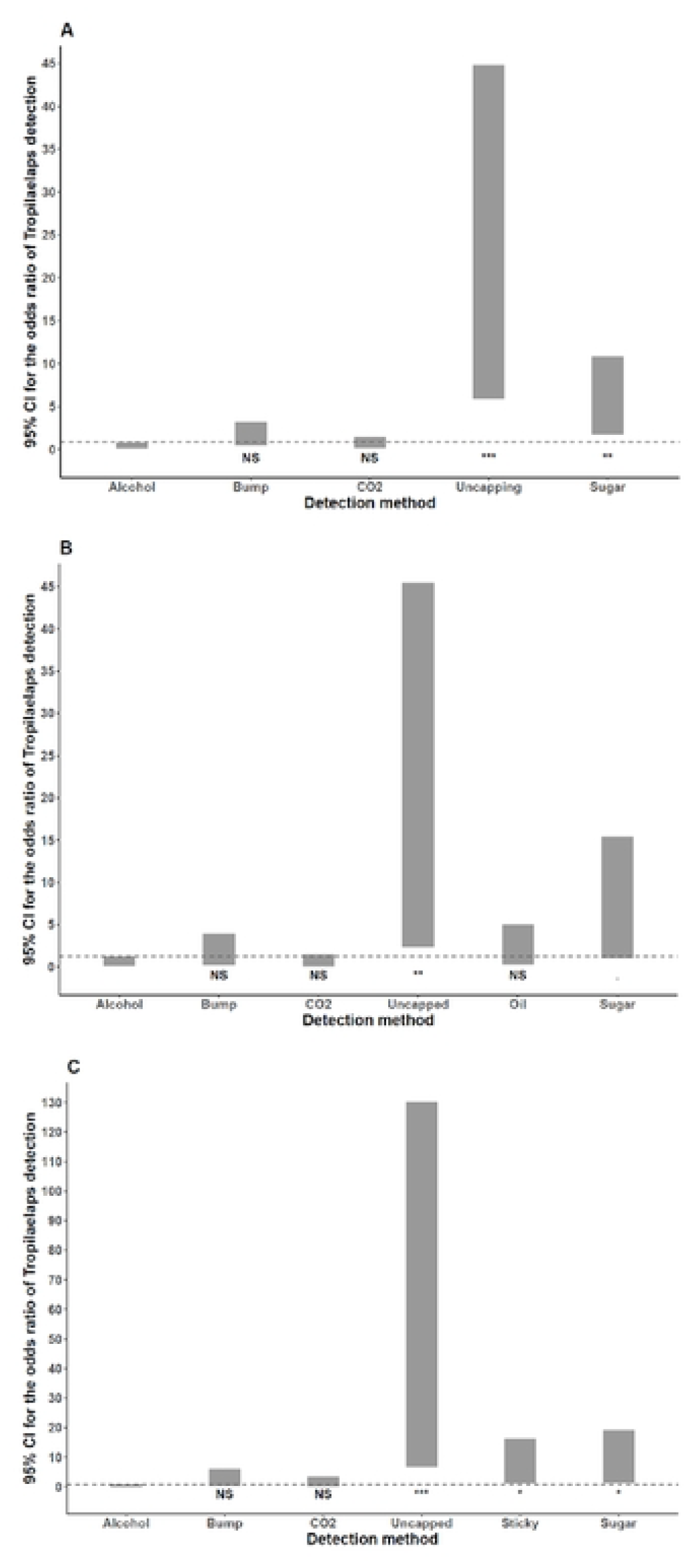
The odds ratio of *Tropilaelaps* detection using different methods when compared to alcohol washing mites from 315 adult bees. The grey bars represent the uncertainty (95% confidence intervals) around the estimated odd ratios. The dashed horizontal line represents the upper 95% CI of the estimate for alcohol. A) All colonies excluding mite drop protocols (n = 53); B) Data restricted to only those colonies where mite drop was monitored using oil (n = 24); C) Data restricted to only those colonies where mite drop was monitored using sticky traps (n = 24).

### Comparing the ability of each method to detect the number of Tropilaelaps mites

Despite efforts to transform the data and to run negative binomial models, the quantitative mite data were not amenable to analysis because of a high number of zeros and then a large span of values for those methods testing positive for mites. The spread of the data are shown (Figure 3) and the raw data are provided (see Source Data).

**Figure 3.**
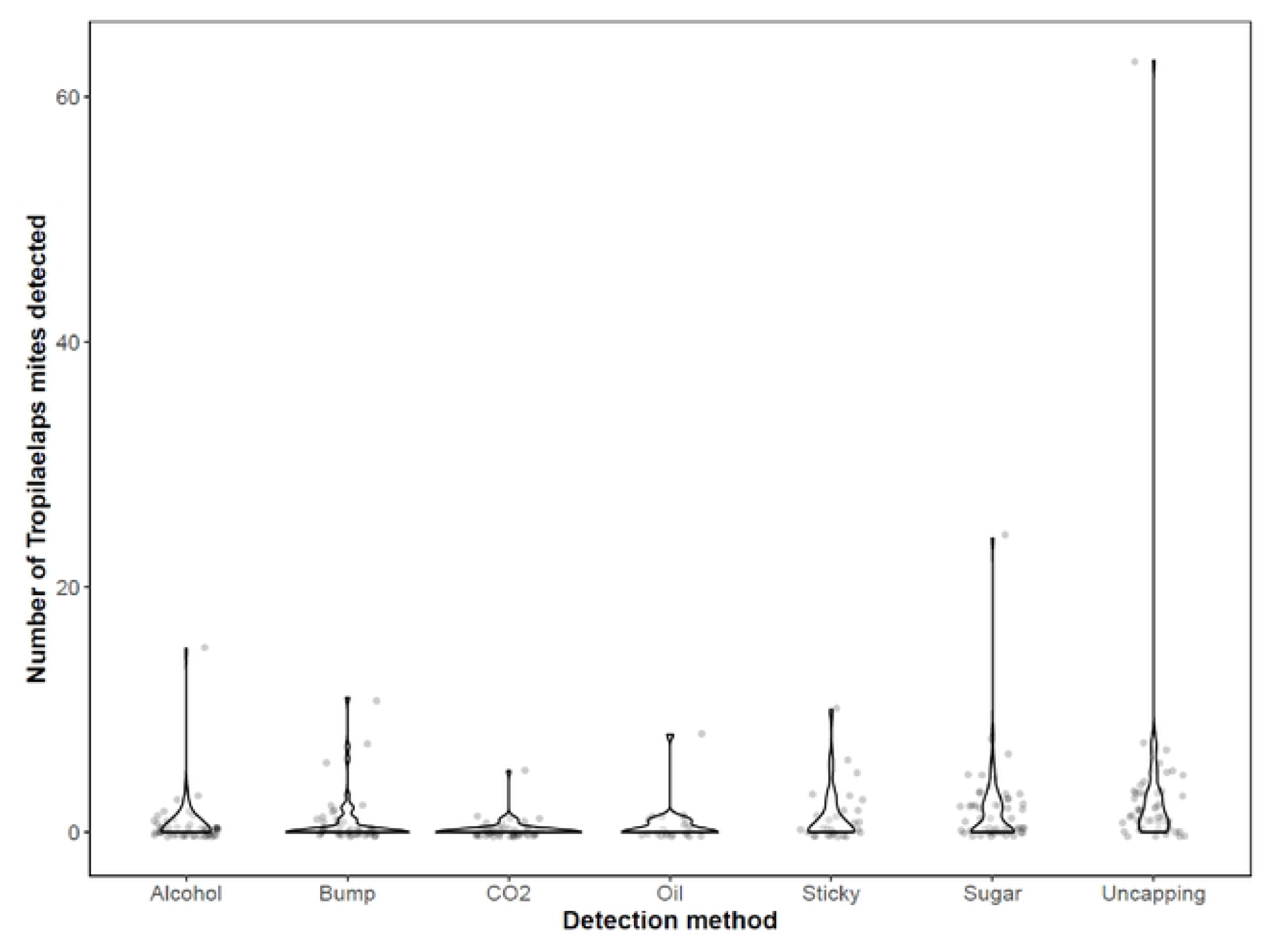
Violin plot of the *Tropilaelaps* count data for all colonies to show all data points and the solid outline highlights the data distribution.

## Discussion

Our study has refined the *Tropilaelaps* mite detection protocols for possible deployment in England and Wales for the purposes of exotic pest surveillance. We established that whilst the use of an uncapping fork is appropriate for the detection of *Varroa* mites, this technique was not suitable for the detection of *Tropilaelaps* due to their size, levels of mobility and fragility. The use of tweezers to remove wax cappings and extract brood was shown to be the most effective detection method for *Tropilaelaps* mites. The selection of post-larval brood was unpredictable as there is no reliable way to determine the developmental stage of brood in sealed cells. Therefore, the technique for assessing if *Tropilaelaps* mites were present in an uncapped cell did need minor adaptation depending on the development stage of the brood removed from each cell. The use of an icing sugar roll of 300 adult bees was also an effective method to detect phoretic *Tropilaelaps* mites in a honey bee colony and was previously untested for the detection of *Tropilaelaps*. An icing sugar roll offers a non-destructive, quick, cheap, and easily deployable detection method that has the potential to be utilised not only during government surveillance but also by beekeepers themselves. Natural mite drop in conjunction with sticky or oiled floors can be reliably used to detect *Tropilaelaps*. Variable light conditions and the need to wear veils made examination of floor inserts in the field an unreliable method for detecting *Tropilaelaps* mites and instead laboratory analysis with good lighting and magnification was needed to conclusively determine if *Tropilaelaps* were present in samples. The screening of floor debris via sieving and ethanol flotation was shown to be completely ineffective for the detection of *Tropilaelaps* mites present in samples. Mites became damaged by the hive debris collection process or entangled in clumps of hive debris making them unrecognisable and the sieving process ineffective. Mites that were dislodged by sieving could not be made to reliably float in 70% ethanol and to achieve flotation the ethanol needed to be diluted to such a point where all the hive debris floated making identification of *Tropilaelaps* mites in a sample impractical. These issues had previously not been identified by other research and this work has resulted in these screening techniques no longer being utilised as part of *Tropilaelaps* surveillance protocols in England and Wales.

An important consideration for anyone assessing the efficacy of field and laboratory methods for the detection of *Tropilaelaps spp* is the level at which a novel infestation needs to be detected to offer the best possible chance for eradication of a pest. Any symptoms of infestation of *Tropilaelaps* mites closely resemble that of *Varroa and* given that beekeepers sometimes struggle to detect *Varroa* mites (Chantawannakul, *et al*. 2018), beekeeper detection seems unlikely. The small size and high mobility of *Tropilaelaps* mites combined with their potential to survive in broodless scenarios increases its potential for transmission around the globe (Truong, *et al*. 2021). Thus proactive, targeted and robust surveillance is crucial to prevent its colonisation of new global areas. Recent novel infestations of *V. destructor* in countries such as La Reunion Island have demonstrated that the introduction of a single mite to a previously *Varroa* free area led to colony losses as high as 64% in some areas during the year following introduction (Esnault, *et al*. 2019). It is reasonable to extrapolate that similar colony losses could be expected if *Tropilaelaps spp*. were introduced to the UK at low levels (Noël, *et al*. 2020) via a similar infection route. Given the limited research and data available on *Tropilaelaps spp*. when compared to other exotic pests such as small hive beetle and *Varroa*, the difficulties associated with detecting *Tropilaelaps*, the similarities in symptomology to a *V. destructor* infestation and the potential high levels of colony mortality a novel colonisation would cause it is crucial that government agencies and beekeepers around the world remain vigilant for this potentially devastating pest.

Our work highlights the importance of testing surveillance procedures for invasive pests in ‘real-life’ inspection scenarios. While previous studies developed detection techniques such as the bump method and brood examination, the practicality and appetite of beekeepers to participate in surveillance using certain techniques needs to be an important consideration. It should not be assumed that surveillance methods will be appropriate for incursions when techniques have been developed in experimental conditions. If the findings of this research are used to advise surveillance procedures, a balance needs to be struck between the use of the significantly more robust detection of *Tropilaelaps* using uncapping with the sacrifice of 100 brood cells versus the non-destructive and more rapid detection of phoretic mites using icing sugar.

Whilst the differences in beekeeping husbandry and climate between the UK and Thailand are significant, we assume these techniques would be easily transferred to the UK. Honey bee behaviour and brood development are comparable between the two countries and factors such as husbandry and climate have limited impact on the *Tropilaelaps / A. mellifera* interactions, as is demonstrated by the ability of *Tropilaelaps* mites to spread to new regions with diverse climates and differing beekeeping practices (Truong, *et al*. 2021).

The use of an alcohol wash or CO_2_ monitoring of adult bees for *Tropilaelaps* mites were rejected as unreliable detection methods. The use of the bump method as part of the standard operating procedure for *Tropilaelaps* mite detection in the UK has now been discontinued based on the results of this study as it was demonstrated to be ineffective and destructive. When colonies were examined 24 hours after the bump method had been used high levels of brood mortality were observed on frames which had been bumped, which had gone unreported in previous similar studies (Figure 4).

**Figure 4.**
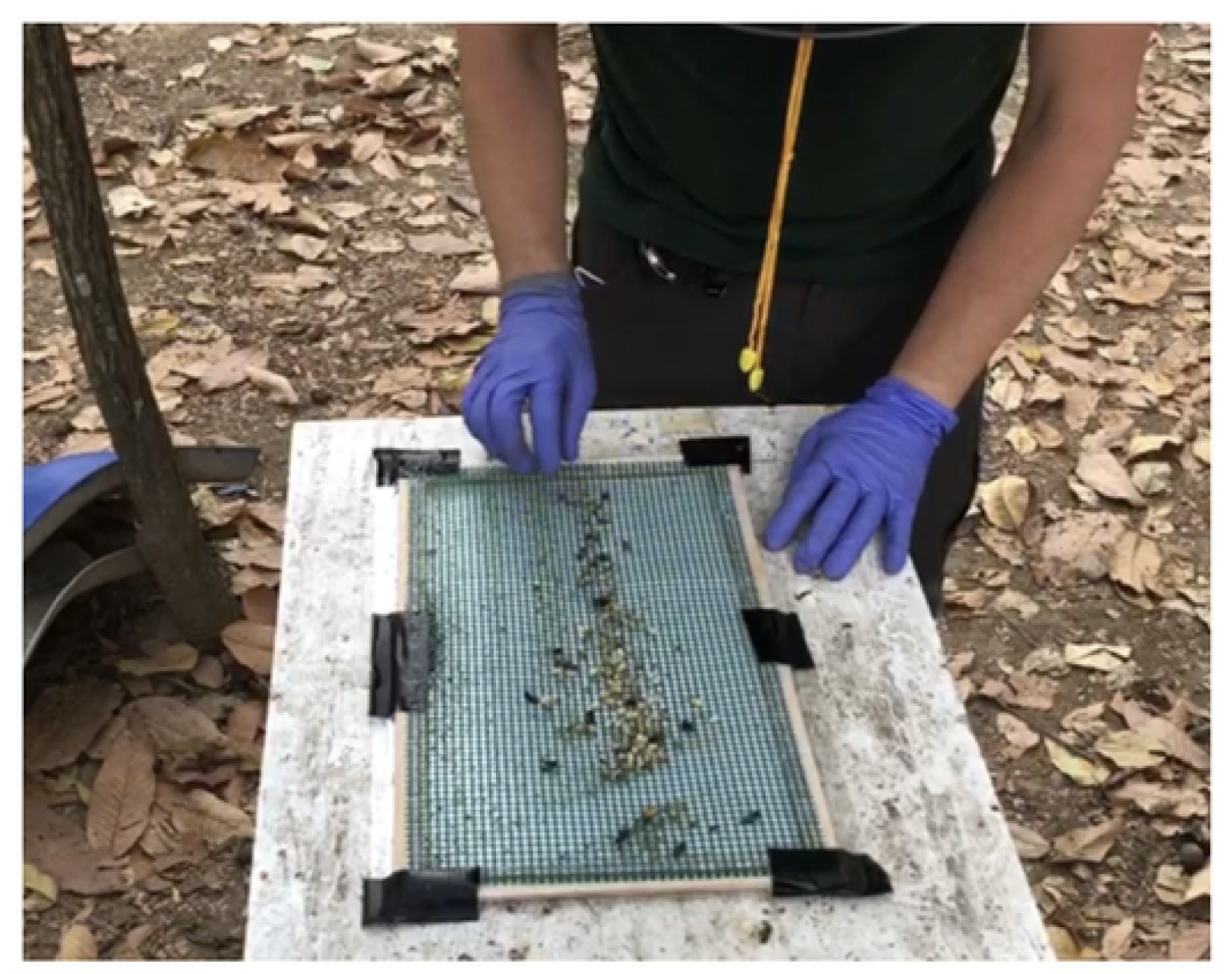
Dead brood on a sticky floor insert placed below a mesh screen from a colony where brood frames had been ‘bumped’ 24 hours previously.

The impact of any surveillance technique on a colony and the practicalities of carrying out any surveillance method in the field in ‘real life’ scenarios should be an important consideration when determining a techniques usefulness. Issues such as brood mortality due to the bump method and the refinement to the brood uncapping technique which were observed during this study where valuable outcomes from this research as determining what doesn’t work in any given situation is as valuable as determining what does. This study has refined and improved existing detection techniques for *Tropilaelaps*, discounted ineffective and damaging detection techniques and given government agencies and beekeepers alike a new, reliable, rapid, non-destructive and cheap surveillance technique. Further work needs to be undertaken to trial the efficacy of commercially available *Varroa* treatments for use during *Tropilaelaps* surveillance and detection (Pettis, *et al*. 2017). Further work is needed to better understand how *T. mercedesae* is able to survive in temperate climates with broodless periods and the implications this might have to beekeeping in the rest of the world should *Tropilaelaps* further spread.

## Acknowledgements

The authors would like to thank Bee Disease Insurance for their generous funding and support of this project. We would also like to thank Dr Bajaree Chuttong and her team at the Meliponini and Apini Research Laboratory at Chiang Mai University for all their assistance.

Giles E Budge was funded by UK Research and Innovation (UKRI) under the UK government’s Horizon Europe funding guarantee [grant number 10082100] as part of the Horizon Europe research and innovation programme BeeGuards [Grant Agreement No. 101082073].

